## Supplementary information for "In vivo self-renewal and expansion of quiescent stem cells from a non-human primate"

**This PDF file includes:**

Supplementary Figures 1 to 4
Legends to Supplementary Figures 1 to 4

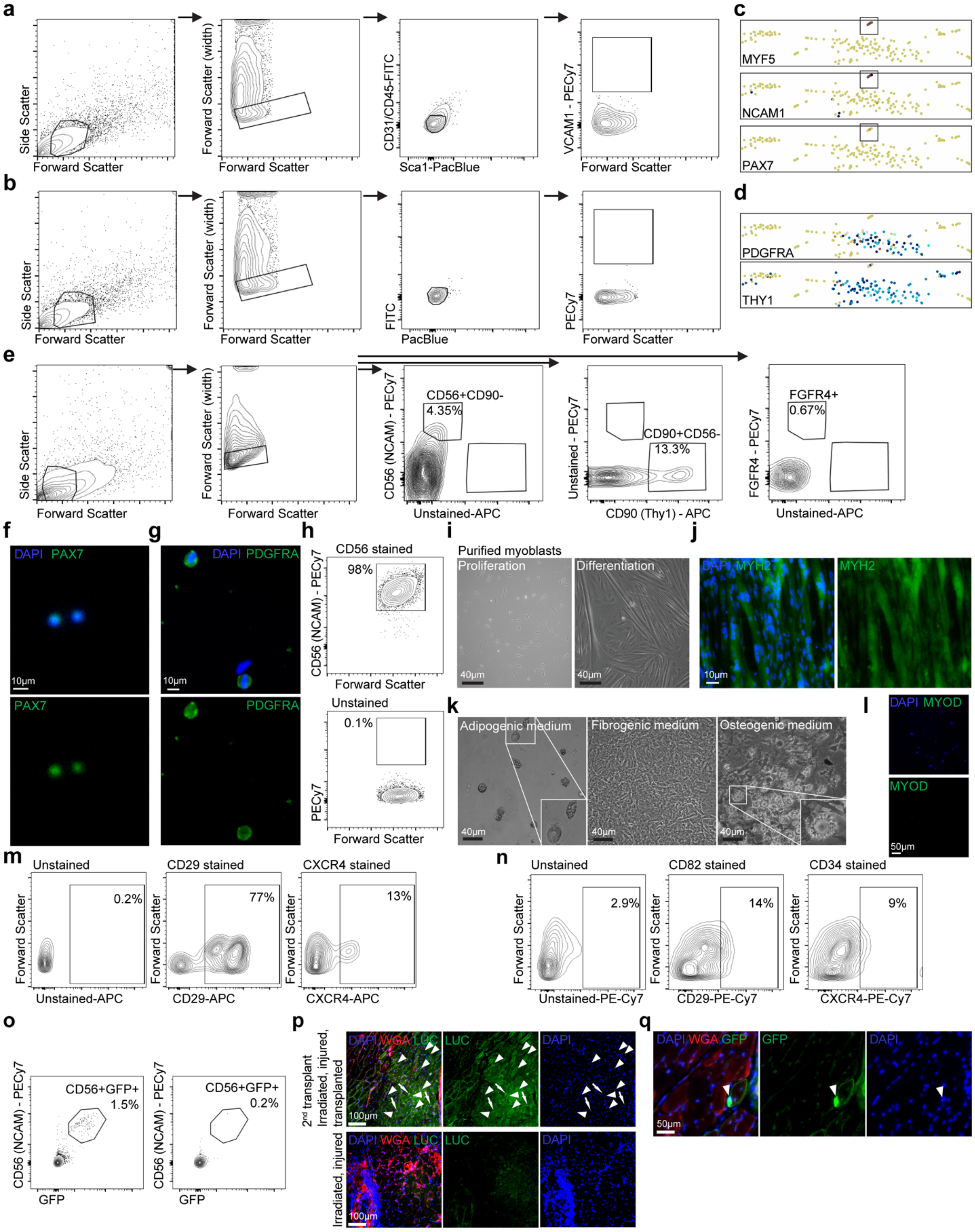


**Supplementary Figure 1: Phenotyping of mouse lemur MuSCs and FAPs.**

(**a, b**) Mouse lemur skeletal muscle tissues were digested into a mononuclear cell suspension and cells were stained with anti-mouse antibody cocktails (a). Cells were subsequently profiled by flow cytometry compared to unstained control (b), first for forward and side scatter (leftmost panels), then by forward scatter width (middle left panels), then by FITC (CD31 and CD45) and PacBlue (Sca1) (middle right panels), and finally by PE-Cy7 (VCAM1) (rightmost panels). (**c, d**) UMAP plots of Smartseq2 single cell RNAseq data of (c) MYF5-positive cells (top panel), NCAM1-positive cells (middle panel), or PAX7-positive cells (bottom panel), and of (d) PDGFRA-positive cells (top panel) or THY1-positive cells (bottom panel), all colored in blue. In (c) the MuSC population is marked by a box. (**e**) FACS plots for anti-CD56 (NCAM1), anti-CD90 (THY1), and anti-FGFR4 single staining controls. Mouse lemur cell suspensions were stained with single antibody clones. Cells were subsequently profiled by flow cytometry, first by forward and side scatter to identify cells, then by forward scatter width to remove doublets, and then by PE-Cy7 (NCAM1), APC (THY1), or FGFR4 staining (panel order from left to right). (**f**) FACS-purified NCAM1$^{+}$THY1$^{-}$ cells stained for PAX7 protein. (**g**) FACS-purified NCAM1$^{-}$THY1$^{+}$ cells stained for PDGFRA protein. (**h**) Myogenic progenitors expanded *in vitro* from purified NCAM1$^{+}$THY1$^{-}$ cells were trypsinized, stained with NCAM1 antibody (right panel), and analyzed by flow cytometry compared to unstained control (left). (**i**) Purified NCAM1$^{+}$THY1$^{-}$ cells were grown in bulk culture in high serum (left panel) or low serum (right panel). (**j**) Myotubes were stained for MYH2 protein. (**k**) NCAM1$^{-}$THY1$^{+}$ cells were expanded for seven days before treatment with indicated differentiation media. Cells were imaged by brightfield. Scale bar = 40 μm. Insets are magnified views of areas noted by the boxes. (**l**) NCAM1$^{-}$THY1$^{+}$ cells were grown in low serum and stained for MyoD protein. (**m**) FACS plots for anti-CD29 and anti-CXCR4 staining. Mouse lemur skeletal muscle tissues were digested into a mononuclear cell suspension and cells were stained with anti-CD29 (middle panel) or anti-CXCR4 (right panel) antibodies and analyzed by flow cytometry compared to unstained control (left panel). (**n**) FACS plots for anti-CD82 and anti-CD34 staining. Mouse lemur skeletal muscle tissues were digested into a mononuclear cell suspension and cells were stained with anti-CD82 (middle panel) or anti-CD34 (right panel) antibodies and analyzed by flow cytometry compared to unstained control (left panel). (**o**) FACS plots of engrafted MuSCs following transplantation. NSG TA muscles that were transplanted with mouse lemur MuSCs were digested and mononuclear cells were stained with an anti-CD56 antibody. Cells were analyzed by flow cytometry for NCAM1 and GFP expression (left). NCAM1$^{+}$GFP$^{+}$ cells were purified by FACS for subsequent analyses. Gates were set to the single cell suspension obtained from untransplanted control muscles, which did not express GFP (right). (**p**) Histology of secondary transplantation from studies described in Fig. 1l. Recipient (2$^{nd}$ transplant) TA muscles or control muscles were fixed and stained with antibodies against luciferase (red) and with WGA to stain the muscle matrix (green). Sections were counterstained with DAPI (blue). Arrowheads mark luciferase-positive central-nucleated myofibers. Arrows mark luciferase-positive mononucleated cells. (**q**) Sections of secondary transplantation were stained for GFP (green) and counterstained with WGA (red) and DAPI (blue). Arrowheads mark GFP-positive cells in the satellite cell position.

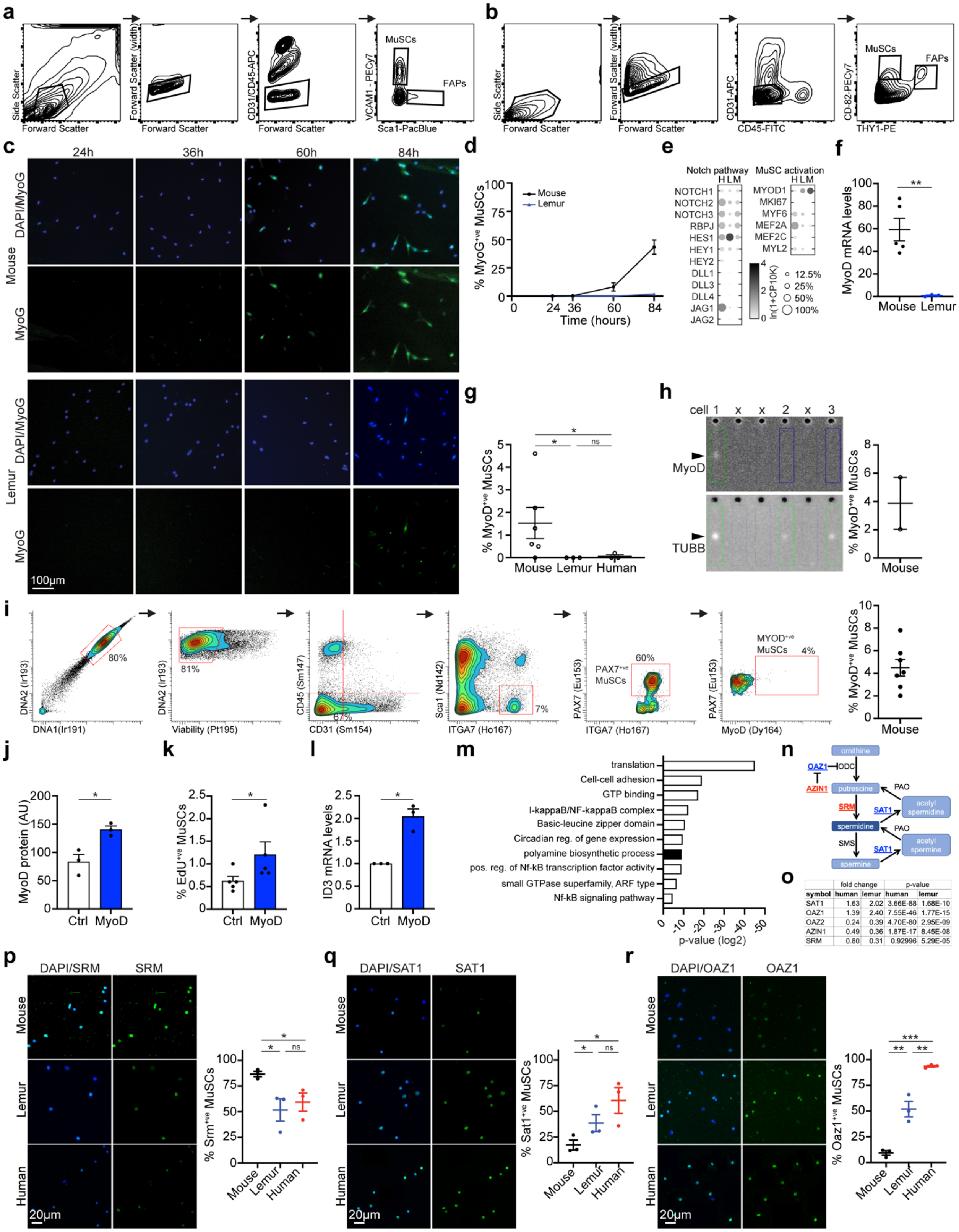


| | fold change | | p-value | |
|---|---|---|---|---|
| symbol | human | lemur | human | lemur |
| SAT1 | 1.63 | 2.02 | 3.66E-88 | 1.68E-10 |
| OAZ1 | 1.39 | 2.40 | 7.55E-46 | 1.77E-15 |
| OAZ2 | 0.24 | 0.39 | 4.70E-80 | 2.95E-09 |
| AZIN1 | 0.49 | 0.36 | 1.87E-17 | 8.45E-08 |
| SRM | 0.80 | 0.31 | 0.92996 | 5.29E-05 |

**Supplementary Figure 2: Altered spermidine metabolism in primate MuSCs.**
(**a,b**) FACS plots demonstrating the gating strategies used to isolate mouse MuSCs and FAPs (a), and human MuSCs and FAPs (b). (**c,d**) MuSCs were grown for indicated times before fixation

and staining for MyoG protein. Representative images shown in (c), graph of quantifications shown in (d). n=3. (**e**) Bubble graph of gene expression levels for canonical NOTCH pathway genes (left) and canonical MuSC activation genes (right) in human (H), mouse lemur (L), and mouse (M) MuSCs (**f**) Microfluidic PCR for MyoD on freshly isolated mouse (n=5 biological independent samples) and mouse lemur MuSCs (n=3). (**g**) Percentage of freshly isolated mouse (n=5), lemur (n=3), and human (n=3) MuSCs that stain positive for MyoD protein. (**h**) Mouse MuSCs were loaded on a single cell western chip and stained for MyoD protein (top left) and beta-Tubulin protein (TUBB, bottom left). The number above the images denotes wells loaded with a cell as called by the TUBB signal. The X above the images denotes wells without signal, in which no cell had been loaded. Lanes containing MyoD$^+$ MuSCs are marked by a red rectangle, while lanes containing MyoD$^-$ MuSCs are marked by a blue rectangle. Graph on the right shows the fraction of MyoD positive MuSCs. n=2. (**i**) Analysis of CyTOF experiments on mouse muscle. Plots demonstrate the gating strategy with the quantification of MyoD$^+$ MuSCs as a fraction of PAX7$^+$ MuSCs on the right. n=7. (**j-k**) Mouse lemur MuSCs were transfected with recombinant MyoD or GFP (Ctrl) for 12 hours and subsequently grown for an additional 24 hours in the presence of EdU. Cells were stained for MyoD protein (j), n=3 biological independent samples, or EdU (k)., n=5. (**l**) Proliferating mouse lemur primary myoblasts were transfected with MyoD or GFP, grown for 24 hours, and assayed for *ID3* mRNA expression. n=3. (**m**) Graph of top fifteen GO term categories that are enriched among the top differentially expressed genes in primate MuSCs. (**n**) Schematic of the spermidine biosynthesis pathway with genes that are expressed higher in primate MuSCs underlined in blue and genes that are expressed higher in mouse MuSCs underlined in red. (**o**) Differentially expressed genes in the spermidine biosynthesis pathway. For each gene, the fold change compared to mouse MuSCs and the adjusted p-value are listed. (**p-r**) Freshly isolated mouse (n=3), lemur (n=3), and human (n=3) MuSCs were stained for SRM (p), SAT1 (q), or OAZ1 (r). Representative images shown on the left, graph of quantifications shown on the right.

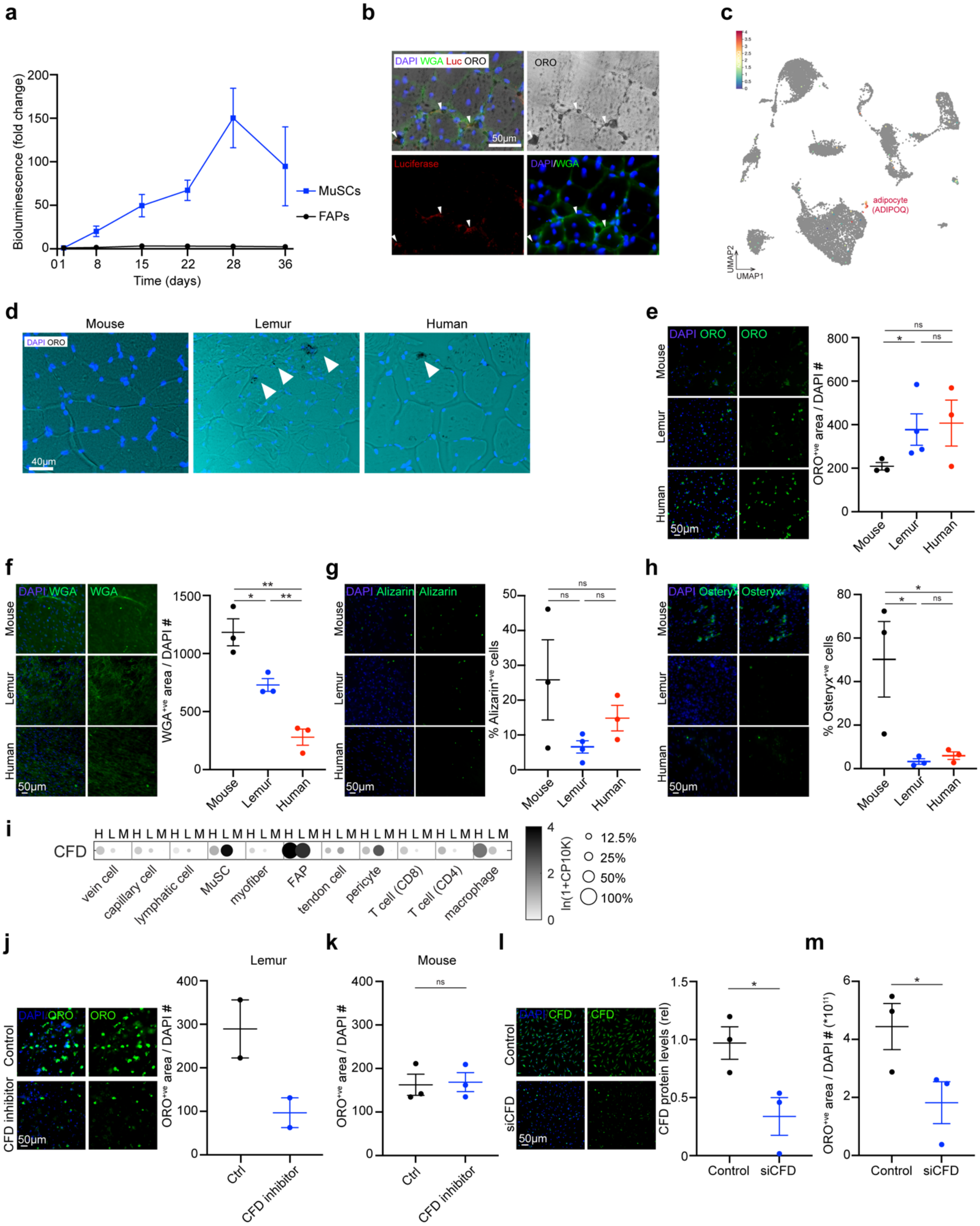


**Supplementary Figure 3: A bias towards adipogenesis in primate FAPs.**
(**a**) Mouse FAPs and MuSCs were transplanted into the TA muscles of NSG mice and engraftment was monitored by BLI. n=12 from 4 donors. (**b**) Mouse lemur FAPs transduced with

luciferase-expressing lentivirus were transplanted into the TA muscles of NSG mice. After one month, muscles were harvested, fixed, and frozen. Cryosections were stained for ORO and with an antibody against Luciferase. Sections were counterstained with DAPI and WGA. Arrowheads denote the interstitial cells that co-stain for Luciferase and ORO. (**c**) UMAP of mouse lemur 10x data with the adipocyte marker ADIPOQ highlighted. n=2 (**d**) Muscle cryosections from mouse (TA muscle), mouse lemur (TA muscle), and human (abdominal muscle) were fixed and stained for ORO (black, red), and counterstained with DAPI (blue). Arrowheads denote the interstitial ORO+ signal present in human and lemur muscle. (**e-h**) Freshly isolated FAPs were expanded *in vitro* and treated with differentiation media for adipogenic (e), fibrogenic (f), or osteogenic (g,h) conversion. Cells were stained for Oil Red O (e), WGA (f), Alizarin Red (g), or SP7 (Osteryx, h). Left panels show representative images, right panels show graphs of quantification. (e,g) mouse n=3, lemur n=4, human n=3; (f,h) n=3. (**i**) Bubble graph depicting gene expression levels for CFD across species and the different cell types isolated from muscle. Species are indicated on the top by their first letter. (**j,k**) FAPs from mouse lemur (i) or mouse (j) were grown in adipogenic medium in the presence of CFD inhibitor Danicopan or vehicle and stained for ORO. The total area of ORO staining was quantified and divided by the total number cells. (j) lemur n=2. (k) mouse n=3. Images of the staining are shown on the left in panel (i). (**l**) Mouse lemur FAPs were transfected with siRNA against CFD or control and stained for CFD. Representative images shown on the left and graph of quantification shown on the right. n=3. (**m**) Mouse lemur FAPs were transfected with siRNA against CFD or control, cultured in adipogenic medium, and stained with ORO. n=3.

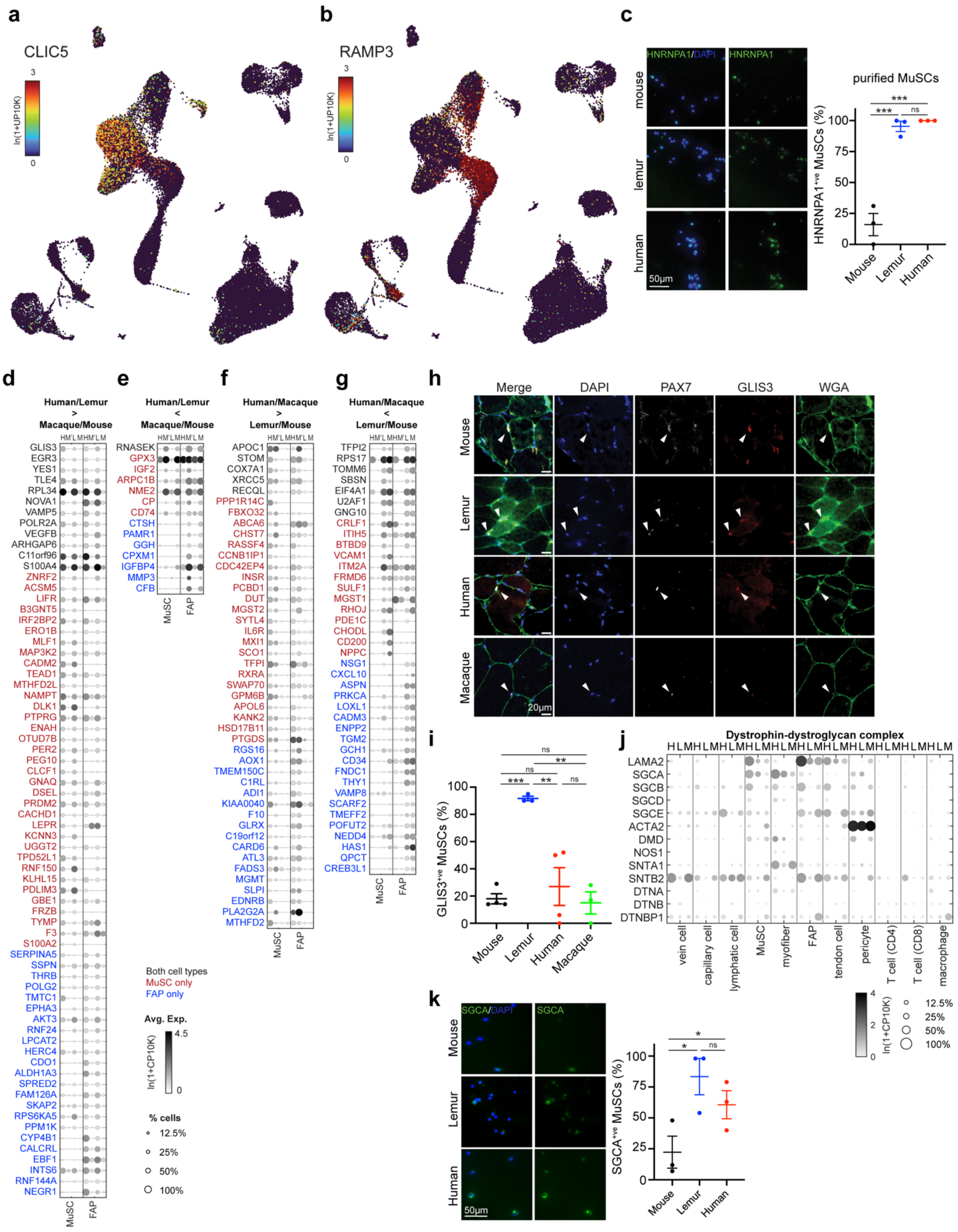


**Supplementary Figure 4: Divergent gene expression in primate stem cells.**

(**a**,**b**) UMAP plot of the endothelial cell compartment of muscle 10X single cell RNAseq data. Colors reflect the expression of CLEC5, a marker for the common capillary endothelial cells (a) or RAMP3, a marker for the capillary endothelial cells unique to macaque (b). (**c**) Freshly isolated mouse (n=3), lemur (n=3), and human (n=3) MuSCs were stained for HNRNPA1 protein and counterstained with DAPI. Images are shown on the left and graph of quantifications is shown on the right. (**d-g**) Stem cell marker plots. Expression levels were calculated and plotted for genes that are higher in the stem cells of two species compared with the two other species. We plotted genes that are higher in human and mouse lemur compared to macaque and mouse (c), genes that are lower in human and mouse lemur compared to macaque and mouse (d), genes that are higher in human and macaque compared to mouse lemur and lemur (e), and genes that are lower in human and macaque compared to mouse lemur and mouse. Cell types are marked at the bottom of each graph. Species are marked at the top of each graph with a single letter: human (H), macaque (M'), mouse lemur (L), and mouse (M). Genes that are significantly different in both MuSCs and FAPs are colored black, genes that are significantly different only in MuSCs are colored red, and genes that are significantly different only in FAPs are colored blue. (**h,i**) Muscle cryosections of indicated species were stained for GLIS3 and PAX7 and counterstained for DAPI and WGA. Arrowheads denote $PAX7^{+}GLIS3^{+}$ cells (h). Graph below in (i) shows quantification of $GLIS3^{+}$ MuSCs for each species. Mouse n=4, lemur n=3, human n=4, macaque n=3. (**j**) Bubble graph of gene expression levels for DGC component genes. Species are indicated on top by their first letter. (**k**) Freshly isolated mouse (n=3), lemur (n=3), and human (n=3) MuSCs were stained for SGCA. A graph of quantifications is shown on the right.
